# Comprehensive annotations of the mutational spectra of SARS-CoV-2 spike protein: a fast and accurate pipeline

**DOI:** 10.1101/2020.06.29.177238

**Authors:** M. Shaminur Rahman, M. Rafiul Islam, M. Nazmul Hoque, A. S. M. Rubayet Ul Alam, Masuda Akther, J. Akter Puspo, Salma Akter, Azraf Anwar, Munawar Sultana, M. Anwar Hossain

## Abstract

In order to explore nonsynonymous mutations and deletions in the spike (S) protein of SARS-CoV-2, we comprehensively analyzed 35,750 complete S protein gene sequences from across six continents and five climate zones around the world, as documented in the GISAID database as of June 24^th^, 2020. Through a custom Python-based pipeline for analyzing mutations, we identified 27,801 (77.77 % of spike sequences) mutated strains compared to Wuhan-Hu-1 strain. 84.40% of these strains had only single amino-acid (aa) substitution mutations, but an outlier strain from Bosnia and Herzegovina (EPI_ISL_463893) was found to possess six aa substitutions. The D614G variant of the major G clade was found to be predominant across circulating strains in all climates. We also identified 988 unique aa substitution mutations distributed across 660 positions within the spike protein, with eleven sites showing high variability – these sites had four types of aa variations at each position. Besides, 17 in-frame deletions at four major regions (three in N-terminal domain and one just downstream of the RBD) may have possible impact on attenuation. Moreover, the mutational frequency differed significantly (p= 0.003, Kruskal–Wallis test) among the SARS-CoV-2 strains worldwide. This study presents a fast and accurate pipeline for identifying nonsynonymous mutations and deletions from large dataset for any particular protein coding sequence and presents this S protein data as representative analysis. By using separate multi-sequence alignment with MAFFT, removing ambiguous sequences and in-frame stop codons, and utilizing pairwise alignment, this method can derive nonsynonymus mutations (Reference:Position:Strain). We believe this will aid in the surveillance of any proteins encoded by SARS-CoV-2, and will prove to be crucial in tracking the ever-increasing variation of many other divergent RNA viruses in the future.

## 1. Introduction

Mutations in the viral genomes serve as the building blocks of viral evolution, and remain the main reason for the novelty in evolution (Baer, 2008; Duffy, 2018). In most cases, mutations are not beneficial for the organisms developing them, and lead to them having fewer descendants over time. Thus, a large portion of mutations, either at nucleotides (nt) and/or change in amino-acids (aa) levels, are harmful (Loewe and Hill, 2010). RNA viruses like SARS-CoV-2 generally have higher mutation rates; in them, however, these mutations are correlated with differential virulence, evolving ability, and traits considered beneficial for viruses (Duffy, 2018; Islam et al., 2020). SARS-CoV-2’s inherently high mutation rate has already produced many descendants from the original Wuhan strain; this complicates its genotyping. The ability of the structural proteins (spike protein especially) in different strains of the SARS-CoV-2 to undergo rapid changes have enabled their genomes to emerge in novel hosts, escape vaccine-induced immunity, and evolve in diverse geo-climatic conditions (Duffy, 2018; Islam et al., 2020; Loewe and Hill, 2010). Moreover, spontaneous mutation is a key parameter in modelling the genetic structure, and evolution of populations (Drake and Holland, 1999). Therefore, investigation of the increased rate of synonymous mutations in the SARS-CoV-2 genomes could be an important tool in assessing the genetic health of populations.

SARS-CoV-2 comprises of four major structural proteins– specifically Spike (S) glycoproteins, envelope (E) proteins, membrane (M) proteins, and nucleocapsid (N) proteins (Ahmed et al., 2020; Rahman et al., 2020; Wu et al., 2020). The entry of SARS-CoV-2 into the host cells is mediated by the transmembrane S protein which consists of two functional subunits responsible for binding to the host cell receptor (S1 subunit), and for fusing the viral and cellular membranes (S2 subunit) (Walls et al., 2020). The higher antigenic and surface exposure properties of the S protein facilitate the attachment and entry of viral particles into the host cells through the host angiotensin-converting enzyme 2 (ACE2) receptor (Grant et al., 2020; Shang et al., 2020; Zhou et al., 2019). Therefore, the spike contains highest variations and determines, to some extent, the viral host range (Coutard et al., 2020; Wu et al., 2020). Furthermore, the S protein is the main target of neutralizing antibodies (Abs) upon infection, and is thus one of the most important structures for therapeutics and vaccine design (Rahman et al., 2020; Walls et al., 2020).

The continuing rapid transmission, and global spread of COVID-19 have raised intriguing questions regarding the evolution and adaptation of SARS-CoV-2 in diverse geographic and climatic conditions driven by synonymous mutations, deletions and/or replacements (Bal et al., 2020; Islam et al., 2020; Pachetti et al., 2020). The capability of the different strains of SARS-CoV-2 strains for swiftly adapting to diverse environments could be linked with their geographic distributions. Though not yet well-studied, evidence suggests that the transmission of SARS-CoV-2 infections and per day mortality rate from this infection is positively associated with weather conditions, and the diurnal temperature range (DTR) (Brassey et al., 2020; Su et al., 2016). However, the exact role of geo-climatic conditions on SARS-CoV-2 is unknown, but it would be worth keeping in mind that this novel disease originated from wildlife before spreading to humans (Harvey, 2020). Therefore, genomic mutation analysis of SARS-CoV-2 strains, integrated with geographic and climatic data, would provide a fuller understanding of the origin, dispersal and dynamics of the evolving SARS-CoV-2 virus. Although several reports predicted possible adaptations at the nucleotide and aa-level, along with structural heterogeneity in viral proteins, especially in the S protein (Armijos◻Jaramillo et al., 2020; Islam et al., 2020; Phan, 2020; Sardar et al., 2020), most of these studies were carried out few complete representative genomes from a limited geographic area. As the genome number is increasing day by day, regular in-house monitoring of the crucial components such as the S protein is urgently necessary to understand the genomic basis and evolution of the diagnostic RT-PCR primer. There are a few pipelines (Yin, 2020) and websites (https://mendel.bii.astar.edu.sg/METHODS/corona/beta/MUTATIONS/hCoV19_Human_2019_WuhanWIV04/hCoV-19_Spike_new_mutations_table.html) in GSAID where aa change or substitution can be observed. In order to provide an alternative tool with a wider range of functions, we present an easy, rapid pipeline that will assist in the alignment of large volumes of viral genomes, remove low quality sequences and in-frame stop codons and provide in-house non-synonymous mutation analysis of large volumes of sequences while requiring minimal knowledge of the command line. This tool can perform this analysis for any other proteins as required. This study aimed to investigate the mutational spectra of aa utilizing this novel methodology in the S proteins in 35,750 complete genome sequences of the SARS-CoV-2 belonging to 135 countries and/regions, and five climatic zones around the world, retrieved from the global initiative on sharing all influenza data (GISAID) (https://www.gisaid.org/) up to June 24, 2020 (Supplementary Data 1).

## 2. Methodology

### 2.1 Genomic data collection, and processing

To decipher the genetic variations of the S glycoprotein, we retrieved 53,981 complete (or near-complete) genome sequences of SARS-CoV-2, available at the global initiative on sharing all influenza data (GISAID) (https://www.gisaid.org/) up to June 24, 2020. These sequences belonged to infected patients from 135 countries and/or regions from across six continents (Supplementary Data 1). Using pyfasta (https://github.com/brentp/pyfasta), we split the total genome into 6 separate files having around 8,900 sequences in each. We aligned each file through the MAFFT (maximum limit 10,000 sequences) online server (https://mafft.cbrc.jp/alignment/server/add_fragments.html?frommanual) using default parameters (Katoh et al., 2002). The complete genome sequence of SARS-CoV-2 Wuhan-Hu-1 strain (Accession NC_045512, Version NC_045512.2) was used as a reference genome.

### 2.2 Mutational frequency analysis

MEGA 7 was used to differentiate the spike protein of SARS-CoV-2 from multiple sequence alignment (Kumar et al., 2016). Sequence cleaner (https://github.com/metageni/Sequence-Cleaner) with set parameters of minimum length (m=3822), percentage N (mn=0), keep_all_duplicates, and remove_ambiguous was employed to remove all ambiguous, and low-quality sequences. We utilized SeqKit toolkit (seqkit grep -s -p “-” in.fa > out.fa) to apprehend gap containing strains for deletion analysis (Shen et al., 2016). Internal stop codon containing sequences were removed by using SEquence DAtaset builder (SEDA; https://www.sing-group.org/seda/). Amino-acid mutation analysis was done with bio-python program using pairwise alignment (https://github.com/SShaminur/Mutation-Analysis). The custom Venn diagrams (http://bioinformatics.psb.ugent.be/webtools/Venn/) server was used to make the Venn diagrams, and visualize the data. Swiss-Model, a structure homology-modelling server (https://swissmodel.expasy.org/) was used to predict the 3D structure (template, PDB ID:6VSB) of the S protein of the reference genome and the structure was visualized in PyMOL (DeLano, 2002; Rahman et al., 2020; Waterhouse et al., 2018). Furthermore, we divided the S glycoprotein mutation of SARS-CoV-2 data according to their geographic origins from six continents - Europe, Asia, North America, South America, Africa, and Australia, and five related climatic zones - temperate, tropical, diverse, dry and continental (Kissler et al.). To estimate the case fatality (mortality) rates of SARS-CoV-2 infections, we collected information on total infected cases, and total reported deaths in these countries from the World Health Organization (WHO) COVID-19 Reports up to June 12, 2020 (WHO Reports, 2020). Microsoft Excel 2016 was used for all the statistical analyses (David, 2017). Detailed step by step methods are described in Mutation_analysis.pdf (https://github.com/SShaminur/Mutation-Analysis).

## 3. Results and discussions

### 3.1 Genomic data collection and processing

Trimming low quality, ambiguous and non-human host RNA sequences resulted in 35,750 (66.23 %) cleaned and full length S protein sequences (Supplementary Data 1). These sequences belonged to 135 countries and/or regions of from six continents (Europe, Asia, North America, South America, Africa, and Australia), and five major climatic zones (temperate, tropical, diverse, dry and continental) around the world (Supplementary Data 1). European countries and/or regions had the highest percentage (58.90%) of S protein sequences, followed by North American (25.78%), Asian (9.34%), Australian (3.61%), South American (1.21%), and African (1.18%) countries or regions. On the other hand, the temperate climatic zone covered the majority of these S protein sequences (60.18%), followed by diverse (33.08%), continental (3.25%), tropical (2.81%), and dry (0.69%) climatic conditions (Supplementary Data 1). We selected the complete genome sequence SARS-CoV-2 Wuhan-Hu-1 strain (Accession NC_045512, Version NC_045512.2) as a reference genome. Through synonymous mutations analysis, we found 27,801 (77.77 %) mutated strains of the SARS-CoV-2 in the cleaned sequences (n= 35,750). Furthermore, country or region-specific aa change patterns revealed the highest number of mutated SARS-CoV-2 strains in England (7,067) followed by USA (6,501), Wales (3,002), Scotland (1,463), Netherlands (1,194), Australia (681), Belgium (596), and Denmark (582) (Supplementary Data 1).

### 3.2 Screening for mutational evolution throughout S protein

Our mutational analyses revealed a total of 988 unique amino acid (aa) change(s)/substitution(s) distributed across 660 unique positions in the S glycoprotein (Supplementary Data 2). The primary structure of the S-protein is 1274 aa, of them 51.81% aa positions (n=660) undergo aa-level evolution worldwide. We found eleven highly variable sites (Position: 32, 142, 146, 215, 261, 477, 529, 570, 622, 778, 791, 1146, 1162) showing four types of aa variations in a single position (Table 1). We also found that positions 52, 185 and 410 in the S glycoprotein had aa variation numbers of 3, 2 and 1, respectively (Fig. 1c, Table 1, Supplementary Data 2). Notably, position 614 showed two variants, substitution D614G (Aspartic acid ◻ Glycine) found in ◻ 74.82 % (n=26,749) of the cleaned sequences (◻96.22% of the mutated sequences), and another variant D614N (Aspartic acid ◻ Asparagine) observed only in four strains from England and Wales (EPI_ISL_439400, EPI_ISL_443658 and EPI_ISL_445498, EPI_ISL_472913). The variant D614G in the S protein has overcome the wild-type variant from China since its first appearance in Germany on January 28, 2020 (Comandatore et al., 2020; Eaaswarkhanth et al., 2020; Kim et al., 2020; Trucchi et al., 2020).

**Table 1.**
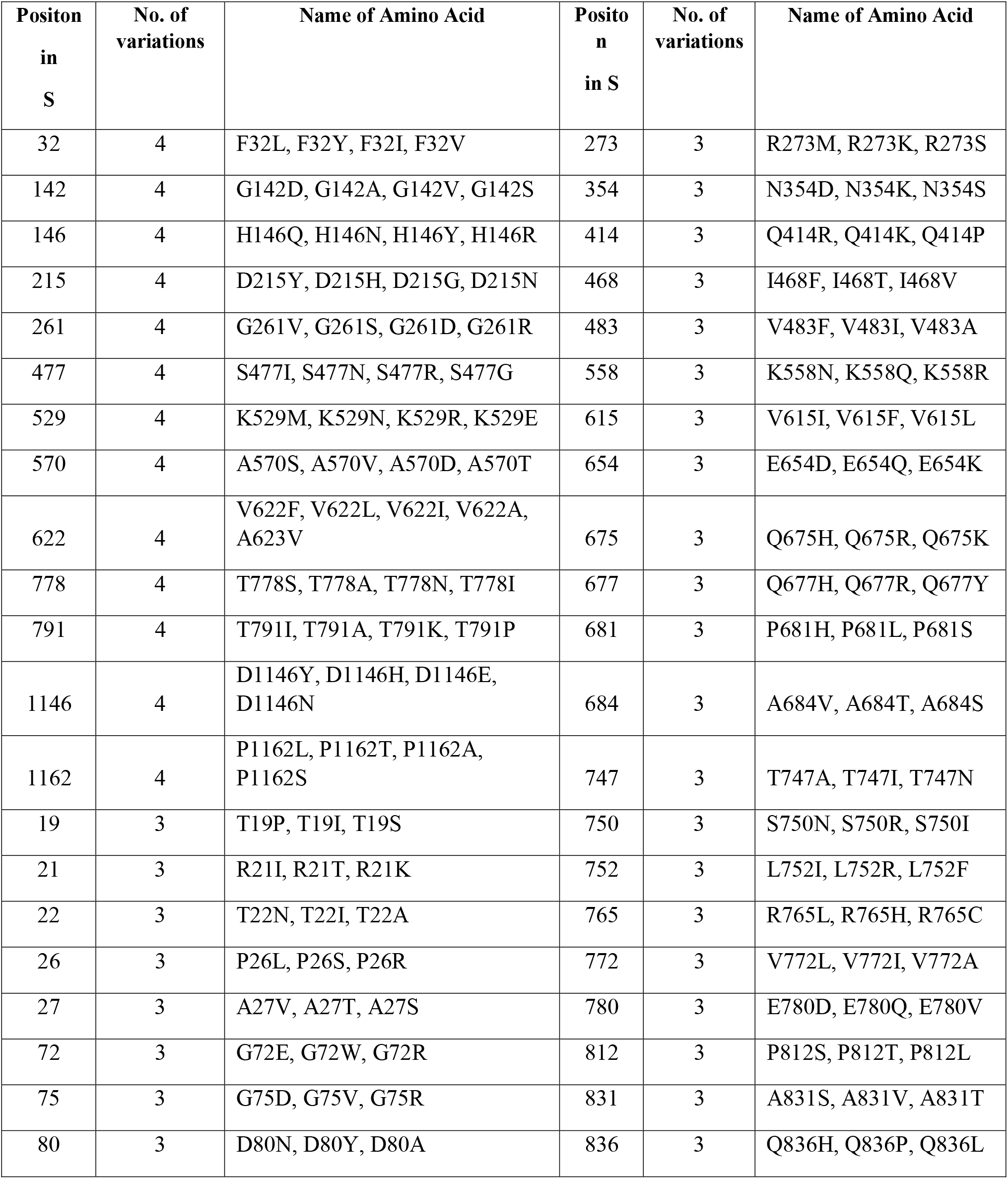
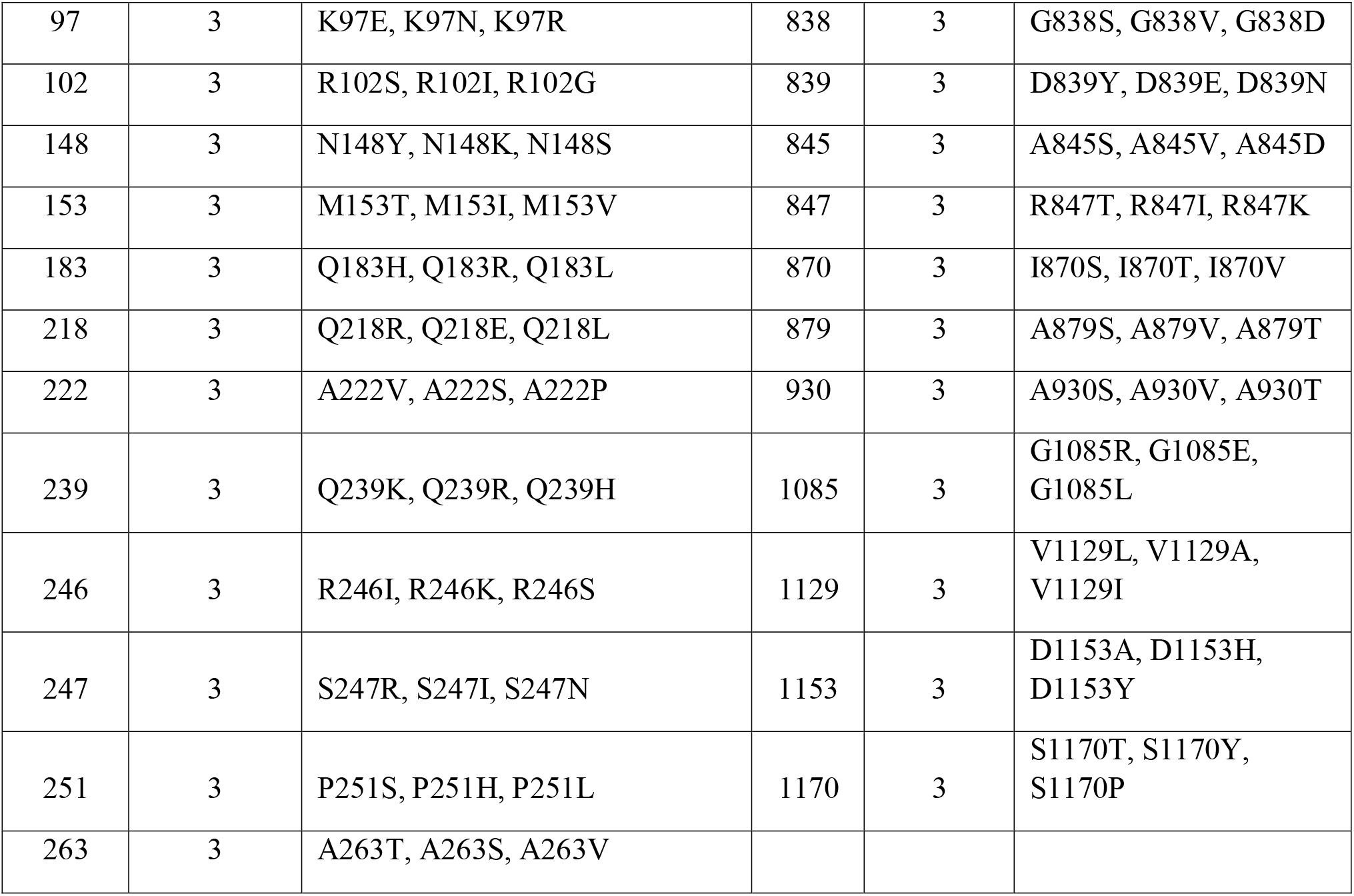
Amino acid variations of S glycoprotein according to their positions. Here, the position where variation more than two aa variations found are represented.

**Fig. 1:**
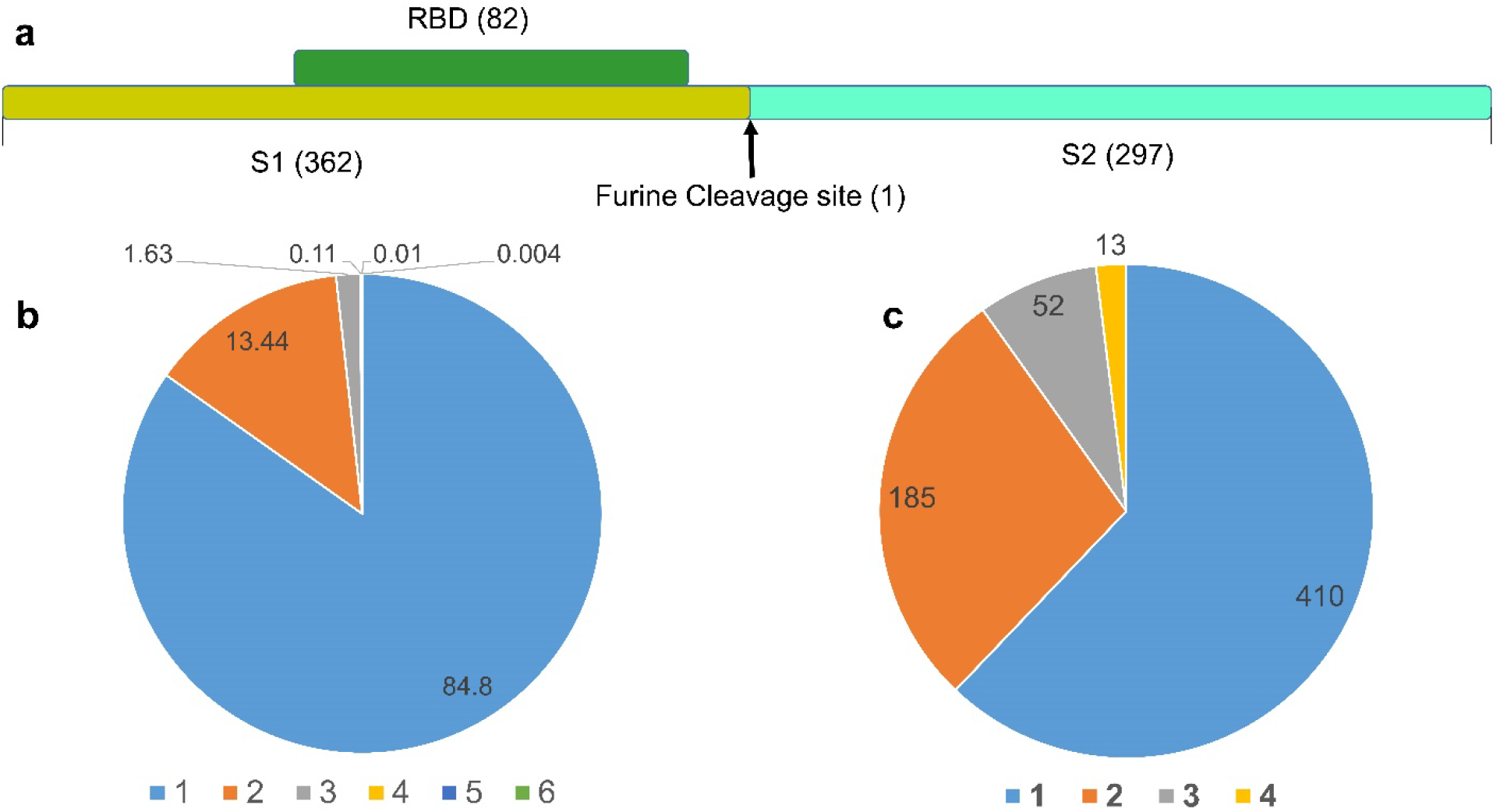
Mutational frequency and distribution of S glycoprotein of SARS-CoV-2. **(a)** Represents different structural regions in spike protein where aa mutations occurred worldwide. The receptor binding domain (RBD) had 82 positions where aa mutations were found whereas the S1 and S2 subunits have 362 and 297 positions for aa mutation, respectively. The furin cleavage site (R685, S686) also possessed one mutation (S686G) in of the Russian SARS-CoV-2 strains (EPI_ISL_428867). **(b)** Denotes the number of mutations in different strains of SARS-CoV-2 where 1, 2, 3, 4, 5 and 6 codes for one, two, three, four, five and six different types of aa mutations across the studied strains. In this study, most of the strains (84.40%) had single aa variation while 13.44%, 1.63%, 0.11% and 0.01% sequences harbored 2, 3, 4 and 5 aa mutations, respectively. **(c)** Positional aa variations in S protein of SARS-CoV-2 where 1, 2, 3 and 4 represent the aa variation in one, two, three and four different positions. 13 positions in the protein were found to having 4 types of aa variations, and 52, 185 and 410 positions in the spike undergone to three, two and one type of aa variations, respectively.

A strain from Bosnia_and_Herzegovina (EPI_ISL_463893) had the highest number of aa changes/substitutions (n=6) at six positions (R246I, L276I, T430A, D614G, S750N, L922V) of S protein. Also, we found that 84.8 % (n=23,576) of the mutated sequences carried just a single aa mutation throughout the S proteins. The remaining 13.44 %, 1.63 %, 0.11 % and 0.01 % of the mutated sequences contained 2, 3, 4 and 5 aa changes, respectively (Fig. 1b, Supplementary Data 2). Moreover, no synonymous mutation was found in the full length S protein of 18 countries and/or regions including Anhui, Brunei, Cambodia, Changzhou, Chongqing, Foshan, Ganzhou, Guam, Hefei, Jiangxi, Jingzhou, Jiujiang, Lishui, Nepal, Philippines, Qatar, Yingtan, Yunnan. This indicates S protein homogeneity of these countries/regions with the reference sequence from Wuhan, China (Supplementary Data1).

The RBD region (Wrapp et., al 2020) (aa position: 338-530) showed nonsynonymous mutations at 82 different positions in 516 strains, whereas in the S1 site and S2 site, there were 362 and 297 positional mutations, respectively. Moreover, in the furin cleavage site (R685 and S686), we also observed a nonsynonymous mutation (S686G) in a single strain (Russia/Krasnodar-63401/2020|EPI_ISL_428867|2020-03-11) (Fig. 1a). We also found aa substitutions at six positions within the RBD region that are directly involved in binding with ACE-2 receptor (Wang et al., 2020; Yuan et al., 2020) including N439K (Scotland, Romania), L455F (England), A475V (USA, Australia), and F456L, Q493L and N501Y (USA) (Supplementary Data 2). All these mutations were found between March and April at a lower frequency (N439K with maximum frequency in 41 Scottish strains and one Romanian strain), except Q493L found in two USA strains reported in May. Q493R position showed variation in an English strain (EPI_ISL_470150) found in April. Furthermore, 18 substitutions at fourteen positions, previously reported to interact with anti-SARS-CoV-2 antibody (Yuan et al., 2020), were found in the strains from Bangladesh, England, Portugal, Wales, Shanghai, France, USA, Scotland, Russia, Latvia, Netherlands, South Africa, Bosnia and Herzegovina, Belgium, Bosnia and Australia (Supplementary Data 2) during the time frame March to May. Discontinuation of the mutants globally may be linked to reduction of virus pathogenicity and virulence fitness affecting transmission dynamics. However, the unavailability of these variants may result due to rejection of the variants with a lower ratio when generating the final consensus sequences as well as insufficient sequences reporting from unusual asymptomatic patients. Moreover, eight glycosylated sites of S protein underwent aa conversions including three substitutions in the NTD region (N17K, N74K, N149H), including a total five substitutions at four sites in the S1 region (N17K, N74K, N149H, N603S, N603K) and four mutations in the S2 region (N717T, N1074D, N1158S, N1194S) (Watanabe et al., 2020). Furthermore, a total of 50 aa substitutions within the S protein observed that incorporated asparagine (N) in S-protein of SARS-CoV-2 including seven within the RBD region (S359N, K378N, K417N, K458N, S477N, T523N and K529N) (Supplementary Data 2). These substitutions alter glycosylation sites and it nature, though it needs further investigations. Overall, the aa substitutions related to asparagine in the RBD (ACE binding domain) and/or in S1/2 domains nearer to the glycosylated sites may affect the glycosylation shield, folding of S protein, host-pathogen interactions, viral entry and finally immune modulation, thus affecting antibody recognition and viral pathogenicity (Ou et al., 2020; Watanabe et al., 2020).

### 3.3 Deletion analysis of SARS-CoV-2 S glycoprotein

Besides site-specific mutations, our analysis revealed 17 in-frame deletions of ranged nucleotides across the SARS-CoV-2 S protein sequences originating from different countries worldwide (Table 2, Supplementary Data 2). Notably, we considered the deletions that occurred in at least two strains at a certain position as deletions. All of the identified deletions distributed throughout the nucleotide sequence 200-2035 fall into four major regions of S protein i.e. nt-positon ranges 179-226 (61-76 aa: NVTWFHAIHVSGTNGT), 413-433 (138-144 aa: DPFFLGVY), 724-732 (241-244: LLAL) and 2021-2035 (675-679 aa: QTQTN). Amino acid deletions at positions 61-76, 138-144, and 241-244 are near the RBD region. Among them, deletions of positions 61-76 and 141-144 are surface exposed, but 241-244 are situated at the inner surface of the predicted S protein (Fig. 2). Also, deleted aa at positions 675-679 are located in the C-terminal transmembrane domain of S protein. Surface exposed deletions near the RBD region may have significant impact on host-pathogen interaction and immune modulation.

**Table 2.**
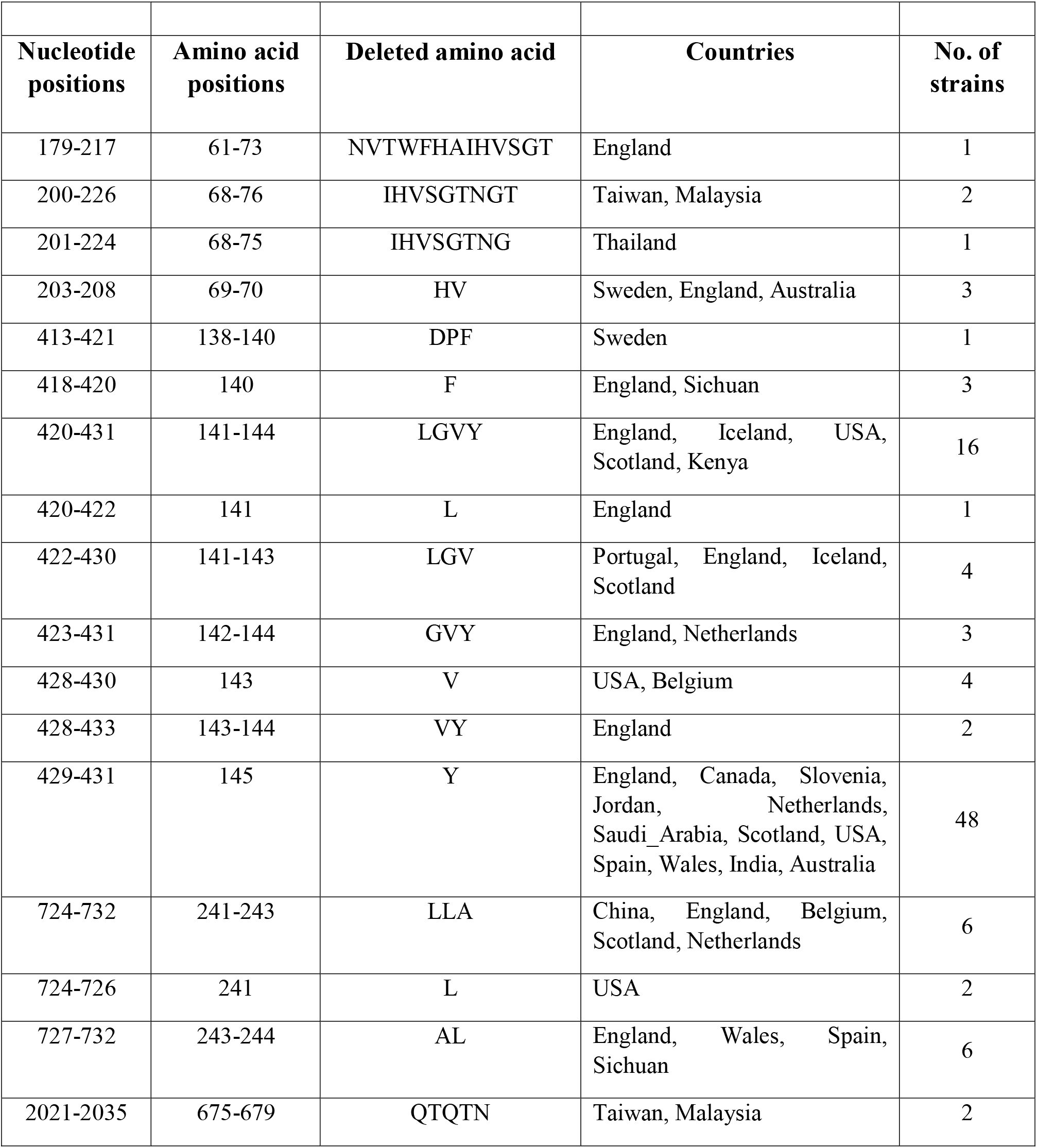
Deletion-sites observed across the S glycoprotein. Countries represent the origin of strains where the deletions found. We considered the deletions that occurred in at least two strains in a certain position.

**Fig. 2:**
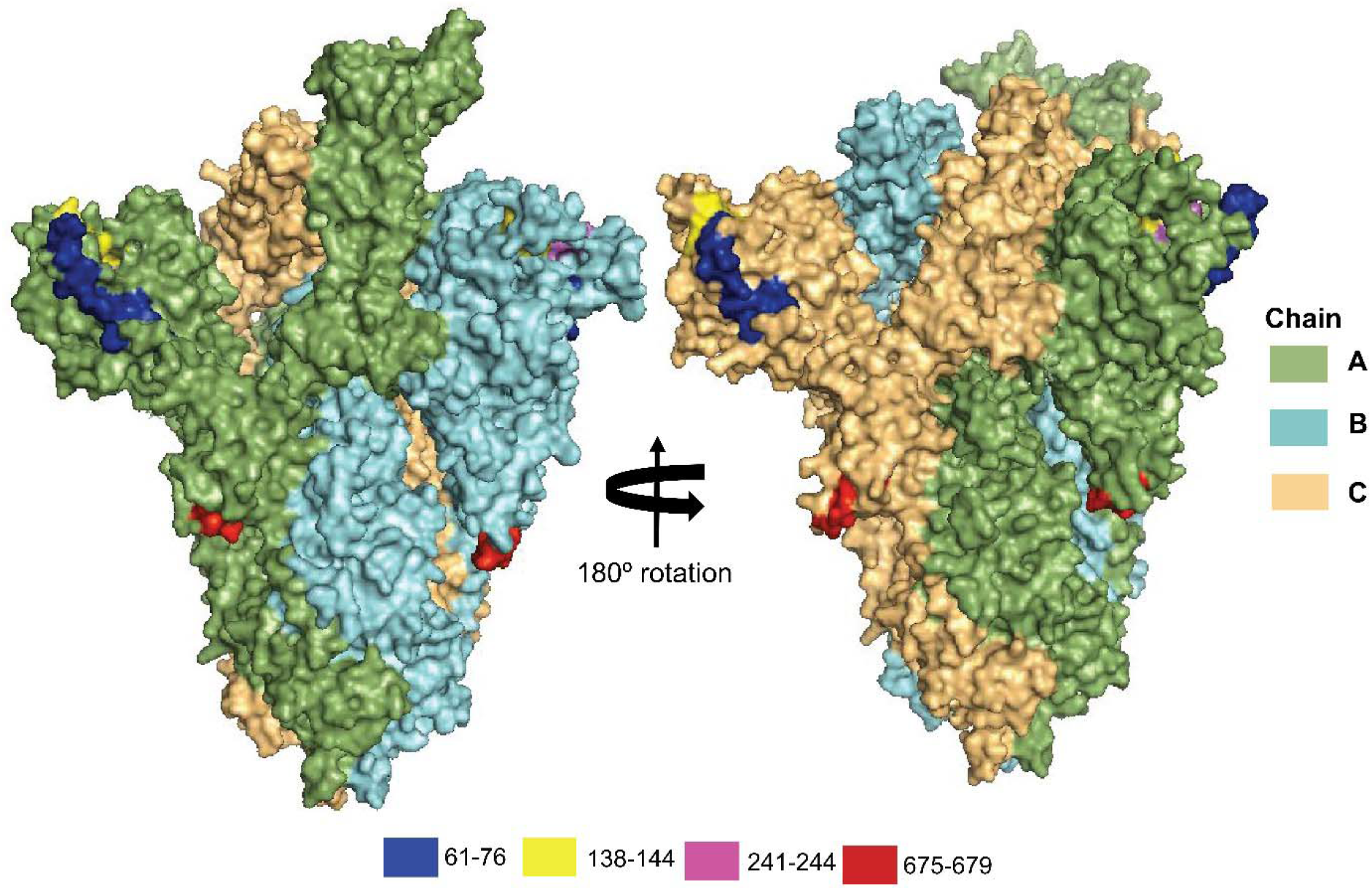
The four amino acids deleted positions (61-76, 138-144, 241-244, and 675-679) in the spike (S) protein of the reference genome, SARS-CoV-2 Wuhan-Hu-1 strain (Accession NC_045512, Version NC_045512.2). The positions are visualized in the tertiary (3D) structure of S protein in PyMOl.

Among the deletions, nucleotide deletion positioned at 418-433 (aa position 140-144) faced frequent overlapped deletions among strains of multiple countries (Table 2). Notably, a single aa in-frame deletion of nucleotides positioned 429-431 (aa position 145) with the highest frequency in 48 strains from multiple countries and/or regions including Australia, England, Canada, Slovenia, Jordan, Netherlands, Saudi_Arabia, Scotland, USA, Spain, Wales and India. A strain from Taiwan (EPI_ISL_444275) showed two coevolving deletions at nt positions 200-226 (68-76 aa:IHVSGTNGT) and nt positions 2021-2035 (675-679 aa:QTQTN). Moreover, two deletions at nt positions 418-420 (140 aa:F) and 727-732 (243-244 aa:AL) were coevolved in a Sichuan strain (EPI_ISL_451369). No other strain had such coevolving deletions, thereby indirectly indicating the negative impact of the deletions on virus fitness and human to human transmissibility. Noteworthy, a 5-aa deletion (675-679 aa: QTQTN) at the upstream of the polybasic cleavage site of S1-S2, and a 21-nt deletion 23596–23617 (aa-NSPRRAR) including the polybasic cleavage site in clinical samples and cell-isolated virus strain likely benefit the SARS-CoV-2 replication or infection in vitro and under strong purification selection in vivo (Liu et al., 2020). Moreover, attenuated SARS-CoV-2 variants with 15-30-bp deletions (Del-mut) at the S1/S2 junction were reported to show less virulence in an animal model (Lau et al., 2020).

These deletions may affect viral adaptations to human, virus-host interactions for infections, attenuation, pathogenicity, and immune-modulations by potentially influencing the tertiary structures and functions of the associated proteins (Phan, 2020). However, further studies are required for the mechanistic clarification and functional implication of these deletions in the SARS-CoV-2 S glycoprotein. The deletion mutations identified in this study should be also considered for current vaccine development.

### 3.4 Geo-climatic scenario of amino-acid changes in the spike protein of SARS-CoV-2, and associated disease severity

Considering geo-climatic impacts on aa changes in the S protein of the SARS-CoV-2, we sought to determine the possible residue positions, and total number of mutations in the S protein gene sequences from 135 countries and/or territories and five climatic zones worldwide. Eight hundred and eighty-eight (988) unique aa replacements across 660 positions along the S protein were identified which differed significantly (p= 0.003, Kruskal–Wallis test) among the genomes of SARS-CoV-2. We found that the frequency of aa changes in the S protein remained substantially higher in the SARS-CoV-2 genome sequences of Europe (62.02%), followed by North America (25.50%), Asia (6.83%), Australia (2.89%), South America (1.41%), and Africa (1.35%) (Supplementary Data 1). Among these replacements, only one aa residue at position 5 (L5F) and 614 (D614G) were found to be the common in Asia, Europe, North America, South America, Africa, and Australia (Fig. 2a). Moreover, 408, 127, 139, 17, 10, and 8 unique aa replacements, and 244, 146, 194, 61, 19, and 23 accessory aa replacements (mutations shared with at least two continents) were found in the SARS-CoV-2 genomes sequenced from Europe, Asia, North America, Australia, South America, and Africa, respectively (Fig. 3a, Supplementary Data 3). Higher unique mutations in European, Asian and American sequences point out the geographical clustering predisposition of the virus. However, further phylogenic study targeting those unique and accessory mutations may lead to a better understanding of global phylodynamics, and thereby guiding the regional control strategy for the COVID-19 pandemic.

**Fig. 3:**
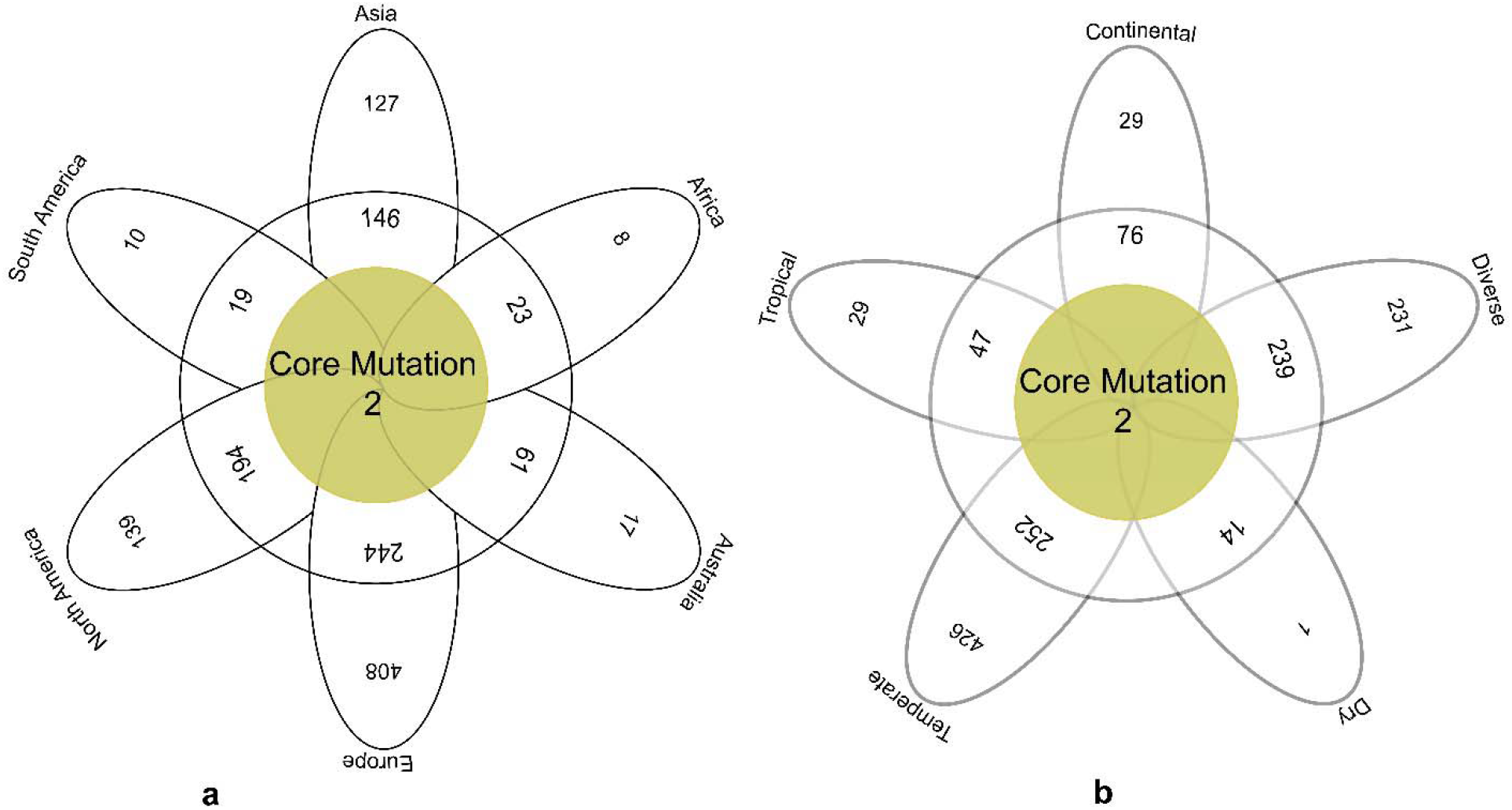
The frequency spectra of amino-acid mutations in the spike protein of SARS-CoV-2. Amino-acid (aa) mutations are represented according to **(a)** geographic areas and **(b)** different climate zones. We found two core shared aa mutation at residue position 5 (L5F) and 614 (D614G) in Asia, Europe, Africa, Australia, North America, and South America, and two core shared mutations at residue positions of 614 (D614G), and 936 (D936Y) in continental, diverse, dry, tropical and temperate climatic conditions. In both cases (a and b), the middle brown circles represent frequency of aa substitutions shared by all variables, and the frequency of aa substitutions shared by at least two continents/climate zones are shown in white circle. The white colored outer ribbons represent unique aa mutations in each individual region and climate zone.

This study also explores the non-synonymous mutations in the S protein of the SARS-CoV-2 genomes across five different climatic conditions worldwide. This revealed significant (p= 0.017, Kruskal–Wallis test) variations in mutation patterns. Our analysis showed that only two core aa substitutions at positions 614 (D614G) and 936 (D936Y) were shared across all the climatic zones (Fig. 3b). Similarly, 426, 231, 29, 29, and 1 unique aa replacement were found in the S protein sequences of the temperate, diverse, tropical, continental and dry climatic conditions, respectively. Moreover, 252, 239, 47, 76, and 14 residue positions in the S protein sequences were identified where nonsynonymous mutations occurred in at least two climatic zones (Fig. 3b, Supplementary Data 3). RNA viruses like SARS-CoV-2 might have remarkable capabilities to adapt to new environments, and confront different selective pressures they encounter (Watanabe et al., 2020).

The genomic variability of SARS-CoV-2 strains manifested by mutations in the spike protein scattered across the globe underly geographically specific etiological effects. One important effect of mapping mutations is the development of antiviral therapies targeting specific regions, for example the spike region of the SARS-CoV-2 genomes (Callaway, 2020). Our current findings corroborate the study completed by Deshwal (2020), who reported the highest SARS-CoV-2 infections and case fatality rates in European countries. In another study, Pachetti et al. (2020) reported two non-synonymous mutations (R203K and L3606F) that were shared across ORFs of the SARS-CoV-2 genomes of six continents, and co-occurrence mutations were also common in different countries along with unique mutations. Nevertheless, mutations in the structural proteins of the SARS-CoV-2, especially in the spike proteins, are driven by the geographic locations that diverged differently, possibly due to the environment, demography, and the low fidelity of reverse transcriptase (Brassey et al., 2020; Pachetti et al., 2020; Su et al., 2016).

Investigating the continental and/or regional impacts of aa substitutions in the SARS-CoV-2 genomes, we found higher case fatality rates in temperate European countries such as United Kingdom (14.16%), Italy (11.72%), France (10.05%), Spain (9.31%), Belgium (3.30%), Germany (3.00%), Russia (2.30%), Netherlands (2.07%), Sweden (1.65%) and Turkey (1.63%) (Supplementary Data 3). Among the tropical Asian countries, higher mortality rates from SARS-CoV-2 infections were estimated in Iran (4.76%), India (4.72%), China (2.56%), Pakistan (1.38%), and Indonesia (1.11%), and rest of the countries had less than 1.0% case fatality rates. Moreover, in the diverse climatic conditions of the American countries or territories (both North and South Americans), United States of America (5.67%), and Brazil (5.14%) had relatively higher mortality rates from SARS-CoV-2 pandemics, and rest of the countries in these continents had substantially lower disease severity rates (< 1.0%). Case fatality or mortality rates from SARS-CoV-2 infections in rest of the two continents (Africa and Australia) remained much lower, and only 2.19%, 1.40%, and 1.26% death rates were found in South Africa, Australia, and Algeria, respectively. The rest of the countries and/or territories of these two continents had less than 1.0% mortality rates (Supplementary Data 3).

The predominantly higher mortality rates in European temperate countries might be correlated with higher unique mutations found in the S proteins reported from this climate Thus, our present study revealed that the predicted rates of unique aa changes in the European sequences could be associated with higher pathogenicity of the virus. However, it is worth noting that reported disease severity (may not represent the actual severity) might be affected by several other factors like health care facilities, average age group and genetic context of the population and control strategies adopted by the countries. Irrespective of the significance of geography for emerging infectious disease epidemiology, the effects of global mobility upon the genetic diversity and molecular evolution of SARS-CoV-2 are under-appreciated and only beginning to be understood. The recent monograph on the spatial epidemiology of COVID-19 makes no reference to the genetic disparity of SARS-CoV-2 (Brassey et al., 2020; Harvey, 2020; Pachetti et al., 2020; Su et al., 2020).

### 3.5 Pipeline validations

The SARS-CoV-2 genomes are increasing very rapidly in the Global initiative on sharing all influenza data (GISAID), but not all genomes are of high quality or complete. So, nonsynonymous mutation analysis with particular crucial part of the virus like S or other structural protein gives statistically more significant insights rather considering the complete genome of the SARS-CoV-2 virus. In this study, we found 33.77 % (18,231/53,981) sequences are in low quality or having ambiguous sequences. Sequence cleaner has removed those sequences and give us cleaned sequences. Among them, we found ten in frame stop codon containing sequences and we have removed it using SEDA. SeqKit toolkit were used to arrest gap containing sequences, and we found around 453 sequences from there, we have to carefully checked the in-frame deletion and 103 strains contains in frame deletions. SNP-sites is a very efficient tools for nucleotide variation detection in different format like multi-fasta alignment, variant call format (VCF), and relaxed phylip format (Page et al., 2016; Seemann, 2015) but this tool is highly dedicated for nucleotide. Snippy (Seemann, 2015) is another tool where nucleotide and protein variation can also be detected, but for large data set with ambiguous sequences will require a separate processing to entrust more accurate results. Here we will get the nonsynonymous mutation that alter aa results in a file format (Sequence_ID Reference_amino_acid:Mutation_Position:Strain_amino_acid) that will assist in the downstream analysis like unique mutation, unique position mutation, mutational frequency, strains having number of mutation. For deletion analysis, this pipeline helps in decreasing the size of sequences (just 453 sequences from 53,981 sequences) for deletion analysis.

## 4. Conclusions

Analyses of genome sequences of 30,493 SARS-CoV-2 strains from across 135 countries and/or regions, and five climatic conditions worldwide revealed the presence of synonymous and non-synonymous mutations, deletions and/or replacements at different positions of the S protein gene, which was reflected in the S-protein primary sequence. These findings of previously unreported mutations in the spike protein of SARS-CoV-2 genomes suggest that the virus is evolving, and European, North American and Asian strains might coexist, each of them characterized by a different mutation patterns, and associated case fatality rates. Moreover, the geo-climatic distribution of the mutations in the spike deciphered higher mutations rates as well as disease severity in the European temperate countries. Furthermore, the structural validations of the mutations in the reference genomes of Wuhan-Hu-1 strain further validated the results of our current study. However, there is no experimental evidence to suggest a difference in aggressiveness of such mutations amongst the studied genome sequences. Moreover, the geo-climate effects of the observed mutations in the spike protein of SARS-CoV-2 on the properties of the diverse strain variants are yet to be evaluated in clinical or experimental studies. Therefore, these results need to be interpreted cautiously given the existing uncertainty about SARS-CoV-2 genomic data to develop potential prophylaxis and mitigation for tackling the pandemic COVID-19 crisis. So, the pipeline developed will help in the easy and accurate way investigate the nonsynonymous mutation, frequency, deletion analysis from large number of data with a shortest possible time without having knowledge of much bioinformatics.

## Conflicts of Interest Statement

The authors of this manuscript declare that they have no conflict of interest.

## Acknowledgements

The authors sincerely appreciate the researchers worldwide who had deposited and shared the complete genomes data of SARS-CoV-2 and other coronaviruses to GISAID (https://www.gisaid.org/). This research utilized these precious data. The authors would also like to extend thanks to Geni Gueiros who was kind to modify his tools (Sequence cleaner) upon request from Md. Shaminur Rahman.

## Data availability

This study utilized the SARS-CoV-2 genome sequences retrieving from the publicly available open database, GISAID. Detailed step by step methods are described in Mutation_analysis.pdf (https://github.com/SShaminur/Mutation-Analysis).

## Author contributions

MSR, MRI, MNH, ASMRUA, MA, JA, and SA conducted the overall study. MSR, MRI, and MNH drafted the manuscript. MNH finally compiled the manuscript. AA, MS, and MAH contributed intellectually to the interpretation and presentation of the results.

## Supplementary Information

Supplementary information supporting the findings of this study are available in this article as Supplementary Files, or from the corresponding author on request.

## Notes

### Competing Interest Statement

The authors have declared no competing interest.

